# Automated quantification of small vessel disease brain changes on MRI predicts cognitive and functional decline

**DOI:** 10.1101/621888

**Authors:** Hanna Jokinen, Juha Koikkalainen, Hanna M. Laakso, Susanna Melkas, Tuomas Nieminen, Antti Brander, Antti Korvenoja, Daniel Rueckert, Frederik Barkhof, Philip Scheltens, Reinhold Schmidt, Franz Fazekas, Sofia Madureira, Ana Verdelho, Anders Wallin, Lars-Olof Wahlund, Gunhild Waldemar, Hugues Chabriat, Michael Hennerici, John O’Brien, Domenico Inzitari, Jyrki Lötjönen, Leonardo Pantoni, Timo Erkinjuntti

## Abstract

**Background and purpose:** Cerebral small vessel disease (SVD) is characterized by a wide range of focal and global brain changes. We used automated MRI segmentation to quantify multiple types of SVD brain changes and examined their individual and combined predictive value on cognitive and functional abilities.

**Methods:** MRI scans of 560 subjects of the Leukoaraiosis and Disability Study (LADIS) were analyzed using automated atlas- and convolutional neural network-based segmentation methods yielding volumetric measures of white matter hyperintensities (WMH), lacunes, cortical infarcts, enlarged perivascular spaces and regional brain atrophy. The subjects were followed up with annual neuropsychological examinations for 3 years and evaluation of instrumental activities of daily living for 7 years.

**Results:** The strongest predictors of cognitive performance and functional outcome over time were total volumes of WMH, grey matter (GM) and hippocampi (p<0.001 for global cognitive function, processing speed, executive functions and memory; and p<0.001 for poor functional outcome). Volumes of lacunes, cortical infarcts and EVPS were significantly associated with part of the outcome measures, but their contribution was weaker. In a multivariable linear mixed model, volumes of WMH, lacunes, GM and hippocampi remained as independent predictors of cognitive impairment. A combined measure of these markers based on z-scores strongly predicted cognitive and functional outcomes (p<0.001) even above the contribution of the individual brain changes.

**Conclusions:** Global burden of SVD-related brain changes as quantified by automated image segmentation is a powerful predictor of long-term cognitive decline and functional disability. A combined measure of WMH, lacunar, GM and hippocampal volumes could be used as an imaging identification model of vascular cognitive impairment.

## INTRODUCTION

Cerebral small vessel disease (SVD) is a frequent cause of stroke and the primary subtype of vascular cognitive impairment. The neuroimaging features of SVD include small subcortical infarcts, lacunes, white matter hyperintensities (WMH), enlarged perivascular spaces (EPVS), microbleeds and brain atrophy,^1^ which have been variably associated with cognitive performance.^2–7^

Recently, SVD has been considered as a dynamic ‘whole-brain’ disease due to the diffuse nature, common microvascular pathologies and varying progression of its lesion types.^8^ A multifactorial approach, taking comprehensive imaging data into consideration with clinical follow-up, may be optimal in characterizing the course of SVD. Visual scoring systems for combinations of MRI findings have been introduced to represent the total load of SVD^9^ and cerebrovascular disease.^10^ These scales are pragmatic but limited in sensitivity. Since the accumulating brain changes in SVD are on a continuum, it is worthwhile to investigate global lesion load with continuous measures rather than ordinal scales.

Automated MRI segmentation has become available for different types of SVD changes. Brain volume, WMH and infarcts have received the most attention, while fewer methods have been presented to quantify lacunes and EPVS. This study aimed to find an optimal combination of the major SVD brain changes by using automated MRI quantification tools. We examined volumetric data of WMH, lacunes, infarcts and EPVS as well as regional brain volume to identify the measures with the strongest associations with cognitive decline. A combined continuous measure was constructed to represent the global burden of SVD brain changes and its predictive value was validated against long-term cognitive and functional outcome.

## METHODS

The data that support the findings of this study are available from the corresponding author upon reasonable request.

### Participants and design

The data was drawn from the Leukoaraiosis and Disability (LADIS) Study, a longitudinal multicenter collaboration investigating the role of age-related WMH in functional disability.^11^ In total, 639 subjects were enrolled at 11 European centers. At baseline, the subjects were 65-84 years old and were classified to have mild to severe WMH on MRI according to the modified Fazekas scale.^11^ They were free of dementia and independent or only minimally impaired in instrumental activities of daily living (IADL) as evaluated with the Lawton IADL scale (score 0-1/8).^12^ The sample represented a mixed population of subjects with either a risk of SVD (at least mild WMH on MRI) or early SVD (more advanced imaging findings, but still independent in daily life).

At baseline, the subjects underwent brain MRI and clinical assessments including a standard neurological examination, functional status evaluation and a neuropsychological examination. In follow-up, the clinical and neuropsychological assessments were repeated annually over 3 years. The number of subjects with clinical evaluation was 639 at baseline, 582 at the 1^st^, 554 at the 2^nd^ and 523 at the 3^rd^ follow-up visit. The number of subjects with neuropsychological assessment was 638, 569, 496, and 468, respectively.

A prolonged follow-up over 7 years was carried out by telephone interviews collecting data from the proxy/informant with the IADL scale focusing on activities in the past 3 months. Conversion from functional independence into disability was defined as an increase of IADL score from 0-1 to ≥2. Of the initial 639 subjects, 94 subjects had died during follow-up. In total, data of outcome in terms of functional disability or death was available for 633 (99%) subjects.

Ethical approval was given by the local ethics committees of each center. Written informed consent was received from all participants.

### MRI acquisition

Brain MRI was administered to the subjects according to the same protocol at each center. The sequences included T1-weighted 3-dimensional magnetization-prepared rapid acquisition gradient-echo, T2-weighted fast-spin echo and fluid-attenuated inversion recovery (FLAIR) images. The initial image analysis was performed centrally at VU University Medical Center, Amsterdam, the Netherlands.^13^

### Image analysis

The automated image analysis methods and their validation results are described in detail in Supplemental material. An overview of the image analysis strategy is presented in Figure 1 and examples of the segmentation results in Figure 2. The types of SVD brain changes were determined on the basis of the STRIVE neuroimaging guidelines.^1^ WMH, lacunes, infarcts and EPVS were automatically segmented using U-shaped convolutional neural networks (CNN). The ground truth segmentations required for training were generated using manual and semi-automatic methods. Volumes of the brain structures were measured from T1 images using an automated image quantification tool (Combinostics Ltd., Finland, www.cneuro.com/cmri/), by which the brain is segmented into 133 regions with a multi-atlas method based on 79 manually segmented atlases (http://www.neuromorphometrics.com/). All volumes were normalized for head size based on a brain size scaling factor.

### Neuropsychological assessment

The LADIS cognitive test battery included the Mini-Mental State Examination (MMSE), the Vascular Dementia Assessment Scale–Cognitive Subscale (VADAS), the Stroop test and the Trail making test.^14^ MMSE and VADAS total scores were used as measures of global cognitive function. Cognitive subdomains of processing speed, executive functions and memory were evaluated with psychometrically robust compound scores detailed in Supplemental material.

### Statistical analyses

The predictive value of each MRI measure on the four cognitive scores in 3-year follow-up was investigated individually with linear mixed modelling (restricted maximum likelihood estimation, unstructured covariance structure) to allow for incomplete data in follow-up. After ruling out multicollinearity, the measures with the strongest associations with cognitive performance in terms of significance of the main effects and MRI*time interactions (indicating a change over time) were entered simultaneously in a multivariable linear mixed model to investigate their independent contributions. The significant measures were then combined as equal components into a single global SVD score by averaging the standardized z-scores of each volumetric measure. Finally, the significance of the individual and combined MRI measures in predicting poor functional outcome was analyzed with Cox proportional hazards models.

All analyses were adjusted for age, sex, years of education and study center. The models with multiple MRI predictors were rerun by additionally controlling for hypertension and diabetes. Since all results remained unchanged, these analyses were not reported. A log transformation was applied for WMH, lacunes, cortical infarcts and EPVS volumes to account for non-normality of the distributions.

## RESULTS

### Sample characteristics

Of the original 639 subjects, 78 cases were excluded from this study because of incomplete set of MRI sequences and 1 case because of failed multi-modal registration in pre-processing. The subjects included in the sample (n=560) did not differ from the excluded subjects in age, sex, years of education, WMH Fazekas score or baseline MMSE score (p>0.05). The characteristics of the sample are shown in table 1.

### Automated MRI segmentation results

The MRI segmentation results of total and regional lesion volumes and brain structures are shown in Supplemental table I and correlations between the main MRI predictors in Supplemental table II.

### Individual associations between the automated MRI measures and cognition

The associations between individual MRI volumes and cognitive scores in 3-year follow-up as investigated with linear mixed models are presented in table 2. After adjustments for age, sex, education and center, all WMH volumes were significantly associated with worse overall performance in all cognitive scores (main effect) and with steeper rate of decline (interaction with time) in processing speed, executive functions and VADAS total score. Total, periventricular and anterior WMH tended to have slightly stronger relationships with cognitive scores compared to the other white matter regions, although the differences between regions were modest.

Total and regional lacunar volumes predicted poorer overall performance in processing speed, executive functions and VADAS, and steeper decline in processing speed, independently of confounders. However, none of the lacunar volumes significantly predicted the memory domain score. Anterior volumes had slightly more prominent effects on cognition than those of the posterior regions.

Cortical infarcts had relatively low predictive value on cognitive performance. Total and regional cortical infarct volumes predicted weakly, although significantly, the overall level and change in processing speed and executive functions. Higher infarct volume was associated with negative main effect on cognitive performance, but there was variability in the estimates of change per follow-up year (data not shown).

Total EPVS volume was significantly associated with overall performance and decline in processing speed as well as decline in memory and VADAS total score.

Total and regional brain volumes showed very strong associations with decline in all cognitive scores. Total cerebral GM and frontal lobe volumes were the strongest predictors of speed and executive functions, whereas hippocampal volume most prominently predicted memory and VADAS total score.

### Independent contributions of the automated MRI measures on cognition

Total volumes of WMH, lacunes, EPVS, cerebral GM and hippocampi were entered together in a linear mixed model adjusted for confounders. Cortical infarcts were left out at this stage because of their weak and variable associations with cognitive decline. As shown in table 3, total WMH volume significantly predicted all four cognitive scores independently of the other lesion types. Lacunar volume independently predicted executive functions. Cerebral GM volume was associated with processing speed, whereas hippocampal volume with VADAS total score, executive functions and memory (table 3). EPVS had no independent contribution to any of the cognitive measures.

### Global score of SVD-related brain changes as a predictor of cognitive functions

A quantitative global SVD measure was constructed by taking the mean of the standardized MRI measures (z scores) significant in the multivariable models above, i.e., total volumes of WMH, lacunes, cerebral GM and hippocampi. The scales of GM and hippocampal volumes were first inversed so that higher scores reflected higher degree of brain changes for all variables. This global score proved to have strong and consistent associations with poorer overall level of performance and steeper rate of decline across all cognitive scores, surpassing the effects of the individual volumes. In linear mixed models identical to individual analyses, the global score significantly predicted processing speed (main effect F 120.6, p<0.001; interaction with time F 37.3, p<0.001), executive functions (F 148.3, p<0.001; F 21.5, p<0.001), memory (F 54.4, p<0.001; F 7.4, p<0.001) and the VADAS total score (F 101.2, p<0.001; F 5.1, p=0.002, respectively).

### Associations between automated MRI measures and functional outcome

Poor functional outcome was defined as subject’s transition to disability or death within the prolonged 7-year follow-up period. As investigated with individual Cox regression analyses controlling for the confounders, poor outcome was significantly predicted by total WMH (HR=4.2, 95% CI 2.9-5.9, p<0.001), lacunar (HR=8.1, 95% CI 3.0-21.7, p<0.001), cerebral GM (HR=0.990, 95% CI 0.986-0.993, p<0.001) and hippocampal volumes (HR=0.57, 95% CI 0.49-0.67, p<0.001), but not by cortical infarct or EPVS volumes (p>0.05). In a multivariable model of the four significant MRI measures, WMH (HR=2.6, 95% CI 1.7-3.8, p<0.001), lacunar (HR=3.2, 95% CI 1.1-9.4, p=0.034) and hippocampal volumes (HR=0.67, 95% CI 0.55-0.81, p<0.001) remained as independent predictors. Finally, the combined score of SVD-related brain changes was significantly related to functional outcome independently of the confounders (HR=2.7, 95% CI 2.2-3.3, p<0.001).

## DISCUSSION

Thus far, there has been no agreed procedure for how to assess the different types of brain changes related to cerebral SVD. In the current study, we leveraged the wealth of information provided by automated CNN- and atlas-based brain MRI segmentation methods to characterize the diverse pathologies of SVD. The volumes with the highest associations with long-term cognitive and functional outcome were identified and the core features were combined to represent the global burden of SVD-related brain changes. The results showed that the combined measure of SVD brain changes was a more powerful predictor of cognitive decline than the individual measures alone.

When considered in isolation, total WMH and GM volumes had the strongest relationships with both global and domain-specific cognitive performance over 3-year follow-up. Hippocampal volume was also strongly associated with cognitive scores, most prominently with global cognitive function and memory domain. Lacunar volumes were significantly related to global cognitive function, processing speed and executive functions, but not with memory. EPVS and cortical infarct volumes had significant, although relatively weak, associations with cognitive decline. In the multivariable models of multiple MRI measures, total WMH, lacunar, GM and hippocampal volumes remained as independent predictors of cognitive functions over 3-year follow-up.

The individual contributions of the automated MRI segmentation volumes are in line with earlier studies using either visual scales or volumetric measures. Within the LADIS study, cognitive impairment has been related to WMH Fazekas score^15^ and semiautomatic volumetry,^4,5^ number and location of lacunes,^4,16^ and visual ratings of global and medial temporal lobe atrophy.^5^ The decisive role of WMH, lacunes and brain atrophy in cognitive decline of SVD has been confirmed by several other studies (see e.g.^3,6,17^). However, despite being a frequent imaging feature of SVD, EPVS have not invariably predicted cognitive decline.^2,3^

We found a close correspondence between the estimates of brain changes detected with the present automated segmentation method and those obtained with manual delineations. Specifically, WMH volume was highly equivalent to the semi-automatic WMH analysis taken as the ground truth.^13^ Lacunes, cortical infarcts and EPVS also reached good accuracy as compared to expert manual delineations (please, see Supplemental material). The imaging data used in this study was relatively heterogeneous multicenter data, and only the 3D T1 had (near)isotropic resolution. Consequently, the imaging data were not optimal but represented the normal clinical variability in the image quality. As the LADIS sample was recruited based on age-related WMH, the number of cases with cortical infarcts was relatively small (in total 73 infarcts) and the training set for the CNN algorithm remained restricted. Cortical infarcts, as a frequent concomitant finding of SVD, showed only a modest predictive value on cognitive impairment, which could be expected given the subcortical predominance of SVD pathology.

The selection of MRI features for the global SVD measure was based on the significance of associations between volumes and multiple longitudinal cognitive and functional outcomes. As the relationship of individual volumes varied between different outcomes, the selected components were given equal weight in the combined score. Importantly, brain atrophy as reflected by reduced global and regional GM volumes was considered together with vascular lesions. Our earlier study has revealed synergistic interactions between atrophy and vascular changes on cognitive decline.^5^ In SVD, WMH and lacunes are closely related to GM atrophy and whole brain volume.^18,19^ In particular, hippocampal and medial temporal lobe atrophy are associated with SVD and contribute to cognitive impairment also in the absence of Alzheimer’s pathology.^20,21^ Brain atrophy is certainly not related to “pure” SVD only, as concomitant neurodegenerative processes are promoted by SVD. In our study, the mechanisms behind hippocampal atrophy remains unclear, as specific biomarkers of concomitant early Alzheimer’s disease were unavailable. More important than to disentangle these processes etiologically, however, is their obvious contribution to predicting cognitive decline and disability.

The current study is one of the first to analyze the clinical significance of automated MRI segmentation features of multiple types of SVD brain changes simultaneously. The visual composite score suggested by Staals et al.^9^ has been found to correlate with general cognitive ability^22^ and decline in executive functions,^23^ but the results have not been quite consistent.^24^ For the visual SVD score, the different lesion types are evaluated using dichotomous ratings.

Continuous volumetric measures are therefore expectedly more sensitive in detecting global SVD changes. Recently, an automated global measure of whole brain atrophy and vascular disease has been proposed and shown to have a stronger relationship with cognition as compared to WMH volume alone and visual total SVD score.^25^ Another study has also reported that a combination of SVD features contributed more to cognitive performance after stroke than the individual measures.^26^ These studies have been limited by relatively small sample sizes and brief cognitive measures. Hippocampal volume has not been included in the measures of global SVD burden before.

Among the strengths of our study are the large and well-characterized sample representing a mixed clinical population of cases with different degrees of SVD, a novel image analysis approach achieving high accuracy in lesion quantification, and the longitudinal design. We applied detailed neuropsychological evaluations to yield psychometrically sound compound indices for both global cognitive function and specific domains. The assessment was repeated annually over three years, while the evaluation of functional outcome in terms of instrumental activities of daily living was extended up to 7 years. The associations were independent of age, sex, education, study center and main vascular risk factors.

Due to unavailable susceptibility weighted MRI sequences we could not include cerebral microbleeds in our analyses, although they are regarded as part of the typical SVD neuroimaging features^1^ and have been identified as risk factors for cognitive dysfunction.^7^ Another limitation of our study is a possible attrition bias in follow-up neuropsychological data (subjects with more severe decline more likely to drop out). Linear mixed models were used as the statistical approach to allow for incomplete observations and utilize all available data.

## CONCLUSIONS

By a robust automated MRI segmentation method capable of identifying multiple types of imaging features of cerebral SVD and brain atrophy, we showed that WMH, lacunar, cerebral GM and hippocampal volumes had the greatest independent predictive value for both cognitive and functional outcome. A combined continuous measure of these four imaging findings was highly predictive of cognitive decline, surpassing the contributions of the individual cerebrovascular disease biomarkers. Global quantification of SVD-related brain changes provides a comprehensive neuroimaging metric of vascular cognitive impairment and may be desirable for SVD intervention studies as a more valid surrogate than single MRI findings.

---

**Table 1.** Characteristics of the study sample (n=560)

|  |  |
| --- | --- |
| Age (mean, SD) | 73.5 (5.1) |
| Sex, women | 303 (54.1%) |
| Education (mean, SD) | 9.7 (3.9) |
| Hypertension | 393 (70.2%) |
| Diabetes | 83 (14.8%) |
| MMSE baseline (mean, SD) | 27.4 (2.4) |
| MMSE baseline <24 | 43/559 (7.7%) |
| MMSE 3-year follow-up (mean, SD) | 26.9 (3.8) |
| MMSE 3-year follow-up <24 | 55/416 (13.2%) |
| Conversion to disability in 7-year follow-up | 281/555 (50.6%) |
| WMH Fazekas score |  |
| Mild | 250 (44.6%) |
| Moderate | 179 (32.0%) |
| Severe | 131 (23.4%) |
| Number of lacunes* |  |
| 0 | 299 (53.4%) |
| 1-3 | 207 (37.0%) |
| $\geq 4$ | 54 (9.6%) |
| Number of cortical infarcts* |  |
| 0 | 515 (92.0%) |
| 1 | 39 (7.0%) |
| $\geq 2$ | 6 (1.0%) |
| Number of enlarged perivascular spaces* |  |
| 0 | 28 (5.0%) |
| 1-5 | 258 (46.1%) |
| 6-10 | 126 (22.5%) |
| $\geq 11$ | 148 (26.4%) |

MMSE indicates Mini-Mental State Examination; WMH, white matter hyperintensities
\* Numbers of lacunes, cortical infarcts and enlarged perivascular spaces were determined with automated MRI segmentation.

**Table 2.** Individual predictive values of automated MRI measures on cognitive functions in 3-year follow-up

|  | **Processing**<br>**speed** | **Executive**<br>**functions** | **Memory** | **VADAS**<br>**total score** |
| --- | --- | --- | --- | --- |
| **White matter hyperintensity volume** |  |  |  |  |
| Total | 64.7 (<0.001)<br>20.7 (<0.001) | 78.2 (<0.001)<br>8.2 (<0.001) | 19.4 (<0.001)<br>2.0 (0.113) | 66.7 (<0.001)<br>5.4 (0.001) |
| Periventricular | 63.9 (<0.001)<br>20.5 (<0.001) | 71.4 (<0.001)<br>6.9 (<0.001) | 31.0 (<0.001)<br>3.3 (0.021) | 74.2 (<0.001)<br>7.1 (<0.001) |
| Deep | 60.8 (<0.001)<br>19.1 (<0.001) | 73.3 (<0.001)<br>8.2 (<0.001) | 13.7 (<0.001)<br>1.4 (0.248) | 57.5 (<0.001)<br>4.6 (0.004) |
| Subcortical | 48.1 (<0.001)<br>17.8 (<0.001) | 46.7 (<0.001)<br>4.4 (0.005) | 10.3 (0.001)<br>1.3 (0.275) | 42.3 (<0.001)<br>3.9 (0.009) |
| Anterior | 67.2 (<0.001)<br>19.2 (<0.001) | 75.7 (<0.001)<br>7.8 (<0.001) | 16.8 (<0.001)<br>1.7 (0.164) | 62.7 (<0.001)<br>5.3 (0.001) |
| Posterior | 53.9 (<0.001)<br>20.8 (<0.001) | 71.0 (<0.001)<br>9.2 (<0.001) | 22.3 (<0.001)<br>2.3 (0.081) | 54.8 (<0.001)<br>5.2 (0.002) |
| Centrum semiovale | 59.1 (<0.001)<br>20.6 (<0.001) | 55.8 (<0.001)<br>7.3 (<0.001) | 12.8 (<0.001)<br>1.4 (0.249) | 36.8 (<0.001)<br>3.4 (0.019) |
| **Lacunar volume** |  |  |  |  |
| Total | 23.8 (<0.001)<br>8.0 (<0.001) | 19.0 (<0.001)<br>1.4 (0.237) | 1.3 (0.256)<br>0.4 (0.748) | 13.3 (<0.001)<br>0.8 (0.474) |
| Anterior | 22.3 (<0.001)<br>7.4 (<0.001) | 16.0 (<0.001)<br>1.1 (0.360) | 1.1 (0.288)<br>0.5 (0.650) | 10.7 (0.001)<br>0.8 (0.494) |
| Posterior | 8.1 (0.005)<br>6.2 (<0.001) | 9.1 (0.003)<br>1.2 (0.316) | 0.3 (0.576)<br>0.4 (0.789) | 8.9 (0.003)<br>0.9 (0.453) |
| **Cortical infarct volume** |  |  |  |  |
| Total | 4.1 (0.045)<br>3.3 (0.020) | 5.3 (0.022)<br>2.0 (0.110) | 0.2 (0.682)<br>0.5 (0.704) | 0.8 (0.375)<br>0.6 (0.616) |
| Anterior | 0.5 (0.462)<br>0.5 (0.667) | 5.0 (0.026)<br>2.3 (0.077) | 0.0 (0.968)<br>0.8 (0.478) | 1.5 (0.222)<br>0.1 (0.980) |
| Posterior | 4.1 (0.042)<br>3.5 (0.016) | 4.4 (0.037)<br>1.6 (0.200) | 0.3 (0.608)<br>0.4 (0.757) | 0.5 (0.487)<br>0.6 (0.598) |
| Right | 7.7 (0.006)<br>5.1 (0.002) | 8.6 (0.004)<br>2.8 (0.037) | 0.1 (0.923)<br>0.6 (0.620) | 3.2 (0.073)<br>0.8 (0.510) |
| Left | 0.2 (0.689) | 0.7 (0.398) | 1.7 (0.199) | 0.1 (0.745) |
|  | 0.5 (0.688) | 0.7 (0.548) | 1.2 (0.309) | 1.0 (0.374) |
| Enlarger perivascular spaces volume |  |  |  |  |
| Total | 4.0 (0.047) | 0.4 (0.530) | 1.7 (0.191) | 1.5 (0.221) |
|  | 6.6 (<0.001) | 1.3 (0.260) | 3.9 (0.009) | 10.6 (<0.001) |
| Regional brain volumes |  |  |  |  |
| Total brain tissue | 49.9 (<0.001) | 74.7 (<0.001) | 45.7 (<0.001) | 56.7 (<0.001) |
|  | 19.0 (<0.001) | 12.4 (<0.001) | 9.0 (<0.001) | 4.2 (0.006) |
| Cerebral grey matter | 103.5 (<0.001) | 99.4 (<0.001) | 33.3 (<0.001) | 45.7 (<0.001) |
|  | 35.9 (<0.001) | 25.8 (<0.001) | 6.3 (<0.001) | 4.1 (0.007) |
| Cerebral white matter | 0.5 (0.485) | 1.9 (0.171) | 8.4 (0.004) | 11.7 (<0.001) |
|  | 2.2 (0.085) | 0.5 (0.672) | 1.4 (0.245) | 1.9 (0.134) |
| Frontal lobes | 101.8 (<0.001) | 98.2 (<0.001) | 23.1 (<0.001) | 40.8 (<0.001) |
|  | 34.9 (<0.001) | 25.0 (<0.001) | 5.3 (0.001) | 3.4 (0.017) |
| Parietal lobes | 76.5 (<0.001) | 68.8 (<0.001) | 27.0 (<0.001) | 27.0 (<0.001) |
|  | 28.9 (<0.001) | 18.5 (<0.001) | 4.4 (0.004) | 4.0 (0.009) |
| Occipital lobes | 66.3 (<0.001) | 49.6 (<0.001) | 3.8 (0.053) | 14.9 (<0.001) |
|  | 36.2 (<0.001) | 11.2 (<0.001) | 0.8 (0.515) | 3.0 (0.032) |
| Temporal lobes | 45.6 (<0.001) | 52.1 (<0.001) | 42.2 (<0.001) | 31.4 (<0.001) |
|  | 11.1 (<0.001) | 15.9 (<0.001) | 11.2 (<0.001) | 2.5 (0.060) |
| Hippocampi | 41.8 (<0.001) | 72.9 (<0.001) | 78.4 (<0.001) | 63.7 (<0.001) |
|  | 15.1 (<0.001) | 14.9 (<0.001) | 11.6 (<0.001) | 3.4 (0.018) |

Linear mixed models adjusted for age, sex, years of education and study center.
Upper row in each cell: Main effect F (p-value) indicates the association between volume and mean cognitive performance across all four evaluations.
Lower row in each cell: Volume\*time interaction F (p-value) indicates the association with change of cognitive performance during follow-up.
VADAS indicates Vascular Dementia Assessment Scale–Cognitive Subscale.

**Table 3.** Combined models for global SVD burden: Independent significance of the MRI predictors on overall cognitive performance over 3-year follow-up

|  | **Processing speed** | **Executive functions** | **Memory** | **VADAS total score** |
| --- | --- | --- | --- | --- |
| WMH | 12.8 (<0.001) | 32.1 (<0.001) | 7.7 (0.006) | 18.4 (<0.001) |
| Lacunes | 0.6 (0.458) | 7.5 (0.006) | 0.0 (0.877) | 2.7 (0.102) |
| EPVS | 0.4 (0.506) | 0.5 (0.469) | 0.1 (0.749) | 0.0 (0.852) |
| Cerebral grey matter | 6.5 (0.011) | 1.8 (0.182) | 0.3 (0.560) | 2.8 (0.096) |
| Hippocampi | 0.1 (0.794) | 4.6 (0.032) | 26.0 (<0.001) | 14.7 (<0.001) |

Linear mixed models F for main effect (p value) adjusted for age, sex, year of education and study center. EPVS indicates enlarged perivascular spaces; VADAS Vascular Dementia Assessment Scale–Cognitive Subscale; WMH white matter hyperintensities.

**Figure 1.**
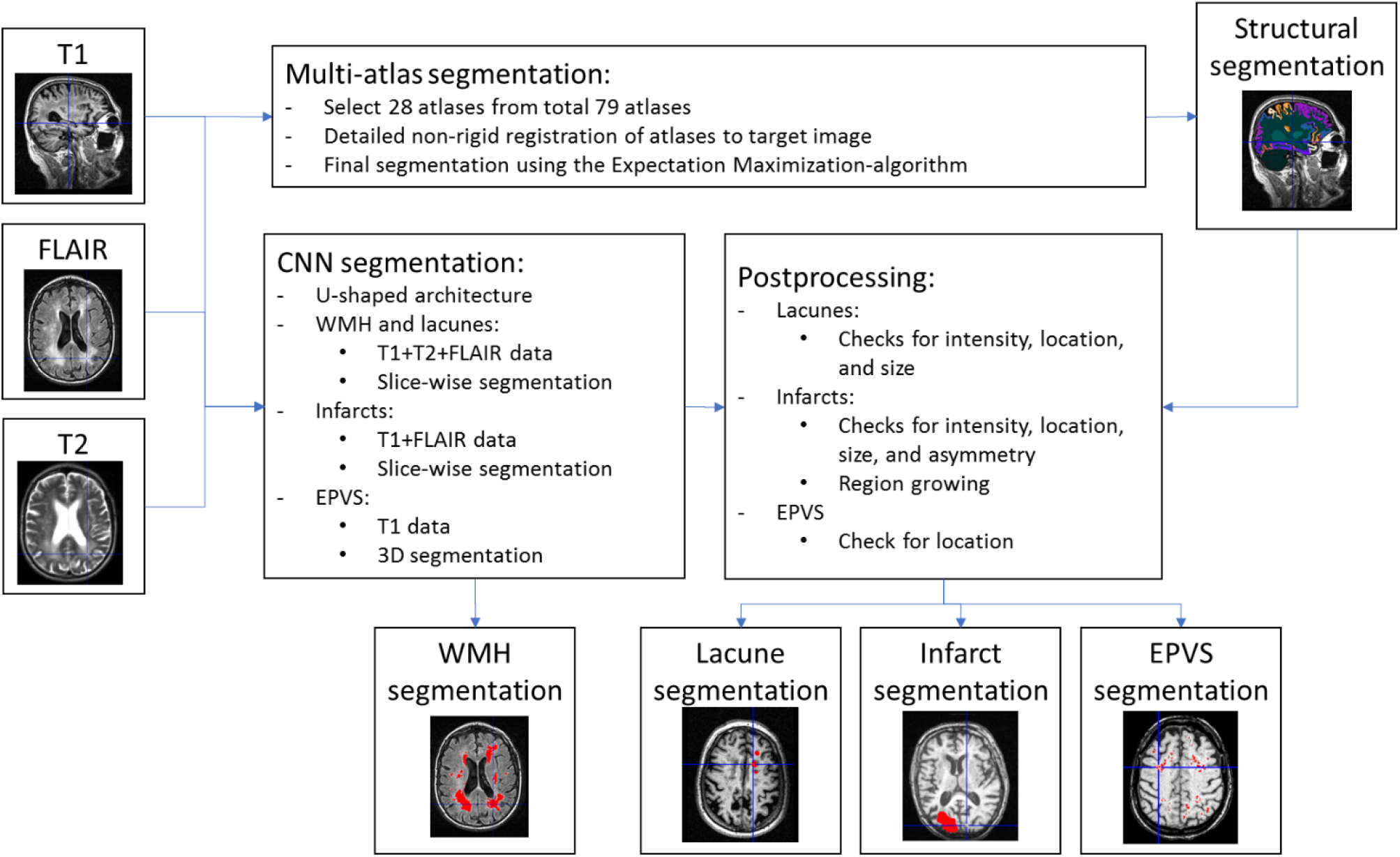
Flowchart of the automated image analysis methods. CNN indicates convolutional neural network; EPVS, enlarged perivascular spaces; FLAIR, fluid-attenuated inversion recovery; WMH, white matter hyperintensities.

**Figure 2.**
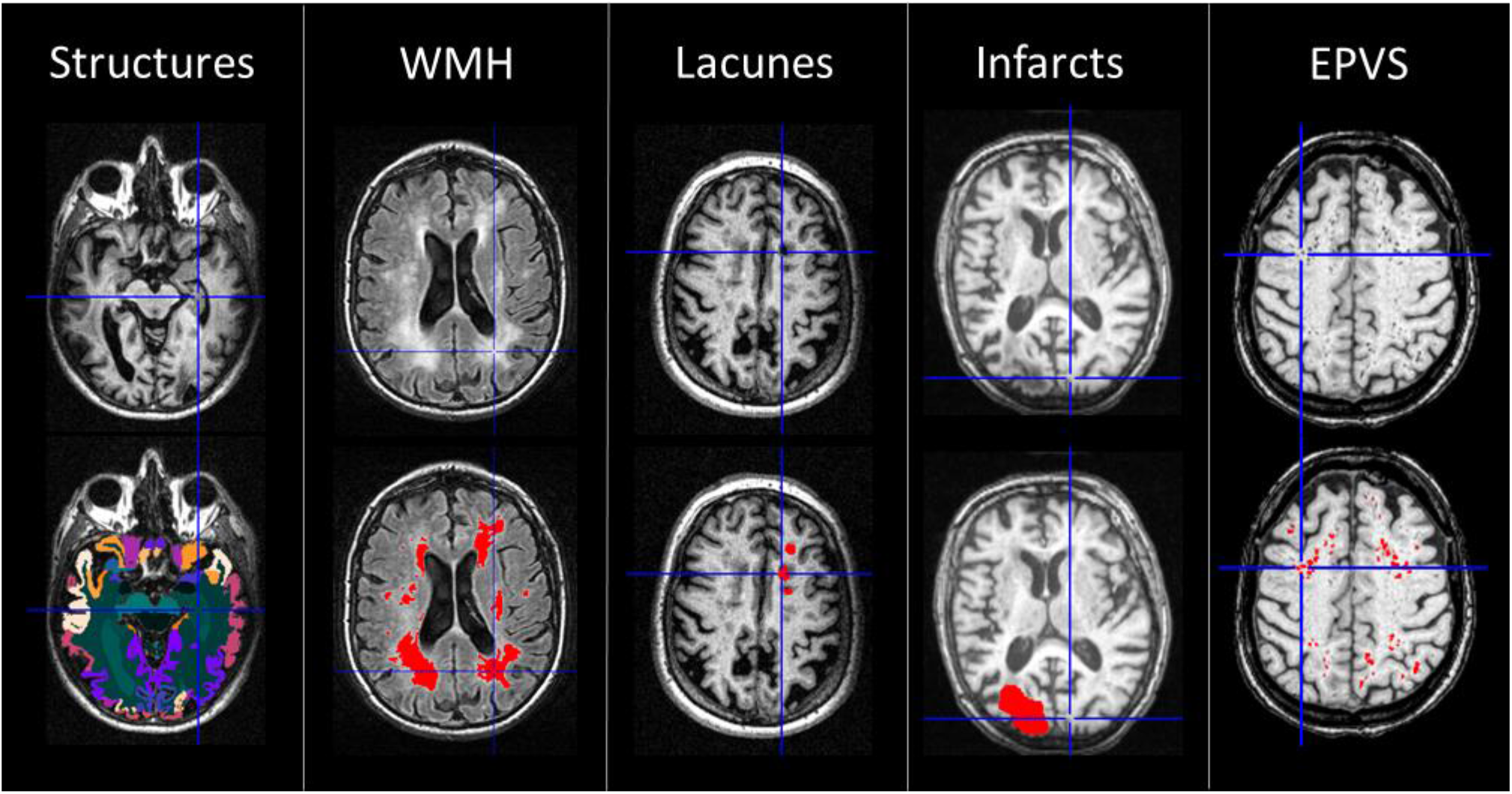
Examples of the automated segmentation results. EPVS indicates enlarged perivascular spaces; WMH, white matter hyperintensities.

