## Supplemental for "Automated quantification of small vessel disease brain changes on MRI predicts cognitive and functional decline"

#### Table of contents

Page

**Supplemental methods**

MRI segmentation method 2

Ground truth segmentations 2

Automated image analysis 2

White matter hyperintensities, lacunes, cortical infarcts and perivascular spaces 2

Volumetry of anatomical regions 3

Head-size normalization 3

Regions of Interest 3

Validation of automated image analysis 4

Neuropsychological evaluation 4

**References** 4

**Supplemental tables**

Table I. Automated MRI segmentation volumes 6

Table II. Correlations among the normalized lesion volumes and structural volumes 7

**Supplemental Appendix**

Leukoaraiosis and Disability Study: List of participating centers and initial personnel 8

### SUPPLEMENTAL METHODS

#### MRI segmentation method

##### Ground truth segmentations

The automated image analysis methods used in this study required ground truth segmentations for training. These ground truth segmentations were generated using manual and semi-automatic methods described below.

The different lesions types of cerebral SVD were determined on the basis of the neuroimaging guidelines of the STandards for ReportIng Vascular changes on nEuroimaging (STRIVE).[1] Hyperintense areas in the white matter on FLAIR sequences without cavitation were regarded as WMH of presumed vascular origin. Lacunes were defined as round or ovoid subcortical fluid-filled cavities (signal similar to CSF) of 3-15 mm in diameter usually having a hyperintense rim in FLAIR sequences. Lacunes were distinguished from EPVS by the difference in size (EPVS generally less than 3 mm in diameter) and shape (EPVS often linear without hyperintense rim). Cortical infarcts were defined as the necrotic tissue located on the cortex (hypointense in T1 and FLAIR sequences).

The ground truth segmentation for WMH was obtained as a part of the initial analysis of the LADIS study, where WMH was semi-automatically defined on the axial FLAIR images in periventricular, subcortical and infratentorial regions.[2] The segmentation was based on a seed-growing technique with local thresholding, where the lesions were marked and borders were set on each slice. Areas of hyperintensity around infarctions and lacunes were omitted.

The ground truth segmentation for lacunes and cortical infarcts were drawn manually from co-registered T1, T2, and FLAIR images using an in-house software tool developed for the manual segmentation of images. All the segmentations were confirmed by an experienced neuroradiologist. In total, there were lacunes in 220 scans (655 lacunes) and cortical infarcts in 55 scans (73 infarcts). The ground truth segmentations for EPVS (53 images, 2759 EPVS) were drawn manually from T1 images using an in-house software tool.

##### Automated image analysis

*White matter hyperintensities, lacunes, cortical infarcts and perivascular spaces*

WMH, lacunes, infarcts and EVPS were automatically segmented using U-shaped convolutional neural networks (CNN).[3,4] CNN are machine learning models that take large number of training samples as an input and build a model that will predict the output based on the training samples. In this study, the training data included T1, FLAIR and T2 images together with the ground truth segmentations and the CNN segmentation was performed using 10-fold cross-validation, i.e., 90% of the dataset was used in training and the rest 10% in testing. This was repeated 10 times so that each image was once used in test set. Three CNN were trained: one for the simultaneous segmentation of WMH and lacunes (utilizing T1, T2, and FLAIR images), one for the segmentation of cortical infarcts (T1 and FLAIR images) and one for the segmentation of EPVS (T1 image). Because only a small portion of the images included lacunes or infarcts, the training of the CNN was performed by emphasizing the training samples with these lesion types. The CNN for EPVS segmentation was trained using the subset with ground truth segmentations. The CNN segmentation for WMH and lacunes and for infarcts was performed slice-by-slice using only 2D data as the slice thickness of the FLAIR and T2 images was typically 5-6 mm, whereas EPVS were segmented using 3D data. The CNN architecture used in this work was the U-shaped residual network presented by Guerrero et al.[3] In short, the network consists of 12 layers with about 1M parameters. There are 8 residual elements, 3 deconvolutional layers, and final convolutional layer that gives the class probabilities for each voxel as an output. The network is trained using image patches (64x64 voxels).

Due to the relatively small training set and large slice thickness, it was noticed that false positive segmentations of lacunes and cortical infarcts occurred. For example, parts of sulci or ventricles could be miss-classified as lacunes in a single 2D slice. Therefore, an automated post-processing step was developed to adjust the segmentations to accurately correspond with the STRIVE guidelines.[1] Multi-atlas segmentation results[5] were used to provide spatial information to remove segmentations from erroneous regions, and a probabilistic atlas of grey-scale values (generated from 534 subjects without major vascular pathologies) was used to provide comparison data for the intensity values. Potential lacunes with diameters <3 mm or missing clear cavity on T1 sequence were discarded. Furthermore, clearly erroneous lacunes located inside ventricles, sulci, or midline were removed. Similarly, potential cortical infarcts that were very small, were not atypically dark in T1, were not located in cortex or did not show asymmetry between hemispheres, were removed. In addition, region growing (dilation five times) was performed to enlarge the cortical infarct segmentations to neighboring voxels that had atypically low intensity in T1. The EPVS segmentations were post-processed by removing possible segmentations located in sulci, ventricles, and lacunes.

*Volumetry of anatomical regions*

Volumes of the brain structures were measured from T1 images using an automated image quantification tool (Combinostics Ltd., Tampere, Finland, www.cneuro.com/cmri/).[6] This tool segments the brain into 133 regions (102 cortical parcellations and 31 sub-cortical regions) using a multi-atlas segmentation method based on 79 manually segmented atlases (http://www.neuromorphometrics.com/).[5] In short, the T1 image of a patient and the atlases are registered using coarse non-rigid deformation. Then, an atlas selection is used to select the 28 best-matching atlases out of the 79 atlases for more detailed non-rigid registration. A probabilistic atlas, generated from these atlas segmentations, is used as a prior in the intensity-based classification using the Expectation-Maximization algorithm.

*Head-size normalization*

All the volumes were normalized for the head size based on a brain size scaling factor derived from the affine registration of loose brain masks of a patient and a reference template.[7]

*Regions of Interest*

In addition to the total WMH/lacune/cortical infarct volumes, following regional volumes were computed: periventricular, deep white matter, subcortical, anterior and posterior, left and right hemisphere, and centrum semiovale. The left-right division and the deep white matter, subcortical, and periventricular regions were defined in the patient space based on the automated multi-atlas segmentation results. The anterior-posterior division and the centrum semiovale were first defined in the Montreal Neurological Institute space (MNI 152-template). The anterior-posterior separation was done using the slice y=110. The centrum semiovale was defined by first extracting the white matter superior to the lateral ventricles (z>32), and then the sulcal white matter regions were removed using a set of morphological operations. The regions of interest were then automatically propagated to the patient images based on a sequence of registrations (MNI – reference template – patient image).

The structural volumetry measures evaluated in this study were the volumes of total brain tissue, cerebral grey matter (GM), cerebral white matter, hippocampi as well as frontal, parietal, occipital and temporal lobes. These volumes were obtained as combinations of the original 133 brain regions of the multi-atlas segmentation.

*Validation of automated image analysis*

Correlation between the WMH volumes of ground truth and automated segmentations was 0.98, and the mean Dice similarity coefficient was 0.77. The corresponding correlation for the volume of lacunes was 0.79, for the volume of cortical infarcts 0.83 and for the volume of EPVS 0.91. The method for the segmentation of structural volumes has been validated in Lötjönen et al.[5] where, for example, the correlation for the volume of hippocampus was 0.94.

#### Neuropsychological evaluation

The cognitive test battery of the LADIS study comprised the Mini-Mental State Examination (MMSE),[8] the Vascular Dementia Assessment Scale–Cognitive Subscale (VADAS),[9] the Stroop test and the Trail making test.[10] Global cognitive function was evaluated with the total scores of MMSE and VADAS.

For the evaluation of cognitive subdomains, three compound measures were constructed by averaging the *z* scores of individual tests within each domain.[11] The scales were first inverted, where necessary, so that higher scores indicated better performance in all variables. Specifically, processing speed was evaluated with the Trail making part A, VADAS Maze task and Digit cancellation. Executive functions were assessed with the Stroop III-II time difference score, Trail making B-A time difference score, VADAS Symbol digit modalities test and Verbal fluency. Memory was evaluated with the VADAS Immediate word recall, Delayed recall, Word recognition and Digit span subtests. The 3-factor model has been supported by confirmatory factor analysis study suggesting that the domains are valid latent variables of cognitive performance and relatively consistent over time.[12]

#### SUPPLEMENTAL TABLES

**Supplemental table I**. Automated MRI segmentation volumes

| | Mean (SD) |
|---|---|
| White matter hyperintensities | |
| Total | 19.3 (19.4) |
| Periventricular | 4.9 (3.2) |
| Deep | 12.2 (14.8) |
| Subcortical | 2.0 (2.4) |
| Anterior | 12.9 (13.6) |
| Posterior | 6.4 (7.1) |
| Centrum semiovale | 2.8 (4.2) |
| Lacunes | |
| Total | 0.22 (0.45) |
| Anterior | 0.19 (0.41) |
| Posterior | 0.03 (0.09) |
| Cortical infarcts | |
| Total | 0.34 (1.79) |
| Anterior | 0.05 (0.62) |
| Posterior | 0.29 (1.56) |
| Right | 0.13 (0.94) |
| Left | 0.05 (0.63) |
| Enlarged perivascular spaces | |
| Total | 0.28 (0.65) |
| Regional brain volumes | |
| Brain tissue, total | 1099.8 (50.0) |
| Cerebral grey matter, total | 483.3 (53.5) |
| Cerebral white matter, total | 415.9 (64.3) |
| Frontal lobe, right | 89.8 (11.2) |
| Frontal lobe, left | 90.4 (11.0) |
| Temporal lobe, right | 53.9 (7.4) |
| Temporal lobe, left | 55.2 (6.7) |
| Parietal lobe, right | 49.5 (6.4) |
| Parietal lobe, left | 49.1 (6.3) |
| Occipital lobe, right | 33.3 (5.5) |
| Occipital lobe, left | 31.5 (5.9) |
| Hippocampus, right | 3.1 (0.5) |
| Hippocampus, left | 3.0 (0.5) |

Ml/brain size scaling factor

**Supplemental table II**. Correlations among the normalized lesion volumes and structural volumes

| | Lacunes | Cortical infarcts | EPVS | Cerebral grey matter | Hippocampus |
|---|---|---|---|---|---|
| WMH | 0.20 (<0.001) | 0.15 (<0.001) | 0.01 (0.798) | -0.41 (<0.001) | -0.42 (<0.001) |
| Lacunes | | 0.08 (0.058) | 0.10 (0.024) | -0.11 (0.012) | -0.03 (0.461) |
| Cortical infarcts | | | -0.1 (0.789) | -0.09 (0.045) | -0.06 (0.161) |
| EPVS | | | | -0.03 (0.449) | 0.00 (0.965) |
| Cerebral grey matter | | | | | 0.67 (<0.001) |

Pearson correlation coefficient (p value)

EPVS indicates enlarged perivascular spaces; WMH, white matter hyperintensities

#### SUPPLEMENTAL APPENDIX

##### Leukoaraiosis and Disability Study: List of participating centers and initial personnel

Helsinki, Finland (Neurology, Helsinki University Hospital and University of Helsinki, Finland): Timo Erkinjuntti, MD, PhD, Tarja Pohjasvaara, MD, PhD, Pia Pihanen, MD, Raija Ylikoski, PhD, Hanna Jokinen, PhD, Meija-Marjut Somerkoski, MPsych, Riitta Mäntylä, MD, PhD, Oili Salonen, MD, PhD; Graz, Austria (Department of Neurology and Department of Radiology, Division of Neuroradiology, Medical University Graz): Franz Fazekas, MD, Reinhold Schmidt, MD, Stefan Ropele, PhD, Brigitte Rous, MD, Katja Petrovic, MagPsychol, Ulrike Garmehi, Alexandra Seewann, MD; Lisboa, Portugal (Serviço de Neurologia, Centro de Estudos Egas Moniz, Hospital de Santa Maria): José M. Ferro, MD, PhD, Ana Verdelho, MD, Sofia Madureira, PsyD, Carla Moleiro, PhD; Amsterdam, The Netherlands (Department of Radiology and Neurology, VU Medical Center): Philip Scheltens, MD, PhD, Ilse van Straaten, MD, Frederik Barkhof, MD, PhD, Alida Gouw, MD, Wiesje van der Flier, PhD; Goteborg, Sweden (Institute of Clinical Neuroscience, Goteborg University): Anders Wallin, MD, PhD, Michael Jonsson, MD, Karin Lind, MD, Arto Nordlund, PsyD, Sindre Rolstad, PsyD, Ingela Isblad, RN; Huddinge, Sweden (Karolinska Institutet, Department of Neurobiology, Care Sciences and Society; Karolinska University Hospital Huddinge): Lars-Olof Wahlund, MD, PhD, Milita Crisby, MD, PhD, Anna Pettersson, RPT, PhD, Kaarina Amberla, PsyD; Paris, France (Department of Neurology, Hopital Lariboisiere): Hugues Chabriat, MD, PhD, Karen Hernandez, psychologist, Annie Kurtz, psychologist, Dominique Hervé, MD, Sarah Benisty, MD, Jean Pierre Guichard, MD; Mannheim, Germany (Department of Neurology, University of Heidelberg, Klinikum Mannheim): Michael Hennerici, MD, Christian Blahak, MD, Hansjorg Baezner, MD, Martin Wiarda, PsyD, Susanne Seip, RN; Copenhagen, Denmark (Memory Disorders Research Group, Department of Neurology, Rigshospitalet, and the Danish Research Center for Magnetic Resonance, Hvidovre Hospital, Copenhagen University Hospitals): Gunhild Waldemar, MD, DMSc, Egill Rostrup, MD, MSc; Charlotte Ryberg, MSc, Tim Dyrby MSc, Olaf B. Paulson, MD, DMSc; Ellen Garde, MD, PhD; Kristian Steen Frederiksen, MD; Newcastle-upon-Tyne, UK (Institute for Ageing and Health, Newcastle University): John O'Brien, DM, Sanjeet Pakrasi, MRCPsych, Mani Krishnan MRCPsych, Andrew Teodorczuk, MRCPsych, Michael Firbank, PhD, Philip English, DCR, Thais Minett, MD, PhD.

The Coordinating center is in Florence, Italy (Department of Neurological and Psychiatric Sciences, University of Florence): Leonardo Pantoni, MD, PhD, Domenico Inzitari, MD (Study Coordinators); Luciano Bartolini, PhD, Anna Maria Basile, MD, PhD, Eliana Magnani, MD, Monica Martini, MD, Mario Mascalchi, MD, PhD, Marco Moretti, MD, Anna Poggesi, MD, PhD, Giovanni Pracucci, MD, Emilia Salvadori, PhD, Michela Simoni, MD.

The LADIS Steering Committee is formed by Domenico Inzitari, MD, Timo Erkinjuntti, MD, PhD, Philip Scheltens, MD, PhD, Marieke Visser, MD, PhD, and Peter Langhorne, MD, BSC, PhD, FRCP who replaced in this role Kjell Asplund, MD, PhD beginning in 2005.
